# The intracellular vacuolar pathogen *Coxiella burnetii* promotes exocytosis in a type IVB secretion system dependent manner

**DOI:** 10.1101/2025.05.26.655767

**Authors:** Hrithik Kumar, Amrita Bhattacharya, Sayanthana Benny, Chandhana Prakash, Nisha Singh, Keerthana Baskaran, Sandhya Ganesan

**Affiliations:** School of Biology, Indian Institute of Science Education and Research (IISER) Thiruvananthapuram, Thiruvananthapuram, India

## Abstract

*Coxiella burnetii* is an obligate intracellular bacterial pathogen that causes a zoonotic disease known as Q fever. Upon internalization into host cells, *Coxiella* extensively remodels host vesicle trafficking and replicates within acidic, lysosome-derived, spacious vacuoles by secreting a suite of effectors through the type IVB secretion system (T4BSS), and hijacking the host cell machinery. However, how fundamental lysosomal functions, such as exocytosis, are subverted during *Coxiella* infection has not been well understood. In this study, we aimed to investigate the regulation and role of exocytosis in the context of *Coxiella* infection. Biochemical and fluorescence-based imaging approaches indicated that *C. burnetii* infection promotes release of extracellular vesicles (EVs). Increase in extracellular levels of endolysosomal proteins (LAMP1 and cathepsin D) and surface LAMP1 expression were identified in both phagocytic and non-phagocytic cells, during later stages of infection. Interestingly, infection-induced exocytosis was dependent on the activity of the bacterial T4BSS. Modulating the activity of TRPML1, a host calcium channel that induces exocytosis by regulating the release of calcium from lysosomes into the cytosol, using synthetic agonist/antagonist, led to a decrease in intracellular *C. burnetii* replication, suggesting a complex, tightly regulated role of TRPML1 during infection. Together, this study demonstrates that infection by the lysosome-adapted pathogen *Coxiella burnetii* activates exocytosis in a temporal, host and bacterial factors-dependent manner.

## Introduction

Lysosomes are highly acidic cellular compartments regulating various functions central to cellular homeostasis, including, but not limited to, catabolism and recycling of nutrients, autophagy, exocytosis, innate immune activation, plasma membrane repair and resealing, membrane remodeling, and antigen presentation (1). Delivery of cellular cargo, as well as microbes to the lysosomes, through multiple vesicle traffic pathways, leads to the degradation of substrates, and this process is a highly conserved cell-intrinsic mechanism to eliminate dysfunctional biomolecules, organelles, as well as pathogens. *Coxiella burnetii*, the causative agent of the zoonotic disease Q fever, is an intracellular bacterial pathogen that obligately replicates inside a host lysosome-derived compartment called the *Coxiella*-containing vacuole (CCV) (2–5). *C. burnetii* uses the specialized Dot/Icm type IVB secretion system (T4BSS) to translocate an array of effector proteins into the host cytosol that subvert host membrane traffic pathways to promote CCV expansion that takes up most of the cell volume over the course of the infection (6–8). The sophisticated adaptation to mammalian intracellular environment, persistent vacuole-bound replication, extensive repertoire of secreted effectors, and subversion of fundamental organelle processes have made *C. burnetii* an excellent model to study organelle biology and the drivers of its dysfunction during infection or other physiologically relevant conditions.

The conversion of a nascent CCV to a spacious, mature, and unified vacuole that supports bacterial replication has been a fascinating, but a less-understood aspect of *C. burnetii* infection. In particular, the mechanisms underlying the potential efflux of toxic build-up and antimicrobial elements from the CCV, over the course of infection, are unknown. Further, it is not well understood whether *Coxiella* intercepts the host exocytic pathway and its contribution to bacterial intracellular replication and dissemination from primary infected cells. In this context, it has been well-established that lysosomes are capable of releasing their content into the extracellular space via lysosomal exocytosis which involves lysosomes fusing with the plasma membrane and also contributing to plasma membrane repair (9–12). Lysosomal exocytosis is regulated by lysosomal calcium channels that facilitate release of calcium from organelle stores to increase intracellular calcium levels, which are subsequently sensed by Ca^2+^ sensor synaptotagmin VII, resulting in fusion of lysosome with the plasma membrane and secretion of lysosomal contents (9,13). Vesicles formed by invagination of late endosomes or intraluminal vesicles of multivesicular bodies can also be released into the extracellular milieu by this process, (called ‘exosomes’) and are considered as important for cell-to-cell communication (14).

One of the key transcription factors that regulates lysosomal exocytosis is the basic helix-loop-helix (bHLH) transcription factor, transcription factor EB (TFEB) (15,16). TFEB is majorly cytoplasmic and is sequestered in a phosphorylated state. The dephosphorylation and nuclear translocation of TFEB is mediated by the lysosomal cation channel, Transient Receptor Potential Mucolipin 1 (TRPML1) (16–18). Release of Ca^2+^ from lysosomes via TRPML1 leads to more cytosolic Ca^2+^, which binds to phosphatases such as calcineurin, making it catalytically active. Calcineurin dephosphorylates TFEB, leading to its nuclear translocation and activation of the transcriptional expression of the Coordinated Lysosomal Expression and Regulation (CLEAR) genes (15,19), including genes involved in lysosomal biogenesis, exocytosis and autophagy (16,17,20). The CCV being a lysosome-derived vacuole, we hypothesized that lysosomal exocytosis may be subverted by *Coxiella* to facilitate periodic extracellular release of lysosomal content to maintain the permissiveness of the CCV for bacterial replication, as well as disseminate infectious bacteria during the course of infection. As the CCV matures and expands in size, by extensively fusing with endolysosomal organelles in the cell, we reasoned that a homeostatic cellular process such as exocytosis may be co-opted to remove/exclude excess organelle load and maintain the CCV permissive for *Coxiella* replication. Interestingly, previous studies strongly implicate TFEB and TRPML1 activity in modulating intracellular *Coxiella* infection, however, finer details on their temporal regulation and specific processes they modulate are required for better understanding of these observations. On one hand, *C. burnetii* infection has been reported to activate TFEB in a T4SS-dependent manner (21) with TFE3/B activity promoting CCV expansion (21,22). On the other hand, TFEB, in particular, has also been demonstrated to negatively impact *Coxiella* replication, with *Coxiella* T4SS activity suppressing agonist-induced TFEB activation (22–24). Further, TRPML1 has been found localized to the CCV, and the *Coxiella* effector CvpE interferes with endogenous TRPML1 activation (25). Based on the above-presented case, we set out to characterize exocytosis during *C. burnetii* infection and identify key host and bacterial factors involved.

## Results

### Infection with *C. burnetii* leads to increased detection of extracellular vesicles (EVs) and endolysosomal proteins in macrophages

EVs are lipid bilayer-enclosed vesicles that carry active and functional biomolecules, including proteins, lipids, nucleic acids, and metabolites, and facilitate cellular homeostasis and cell-to-cell communication (26). Multiple pathways govern EV production and secretion. Some of the major types of EV include (i) exosomes, released by fusion of late endosomal (LE) compartments or multivesicular bodies (MVBs) with the plasma membrane (PM), thereby releasing luminal contents and intraluminal vesicles (ILVs), (average size 30-200nm) (ii) microvesicles (also ectosomes), that are generated by outward budding of the plasma membrane (average size 100-1000nm), (iii) plasma membrane blebbing or fragmentation to generate apoptotic bodies (average size >1000nm), (iv) blebbisomes, that are large membrane bound, nucleus free vesicles containing subcellular organelles (average size ∼10um) (27–30). Thus, EVs comprise distinct populations, are released by virtually all cell types, and are defined by their biogenesis pathways, size and composition.

To investigate potential infection-induced exocytosis and release of endolysosome-derived vesicles and cargo, we infected the J774 macrophage cell line with GFP-expressing *C. burnetii* at a multiplicity of infection (MOI) of 50 for 5 days. As per the schematic shown in **Figure 1A**, the cell-free supernatants were collected on 2 dpi or 5 dpi and first centrifuged at 400g, 10 mins to remove the floating/dead cells, the resulting supernatants were further spun at 3220g, 20 mins to collect extracellular *Coxiella* and the final supernatants were further sequentially centrifuged at 100,000g for 2 hrs to isolate EVs. These fractions were saved and are referred to as “3220g and 100,000g fraction (or EV fraction)” respectively in the rest of the sections. These fractions were immunoblotted to examine the intra- and extracellular abundance of specific proteins that originate from distinct subcellular compartments **(Figure 1B and 1C)**. The absence of nuclear protein lamin A/C in the 3220g and the EV fraction suggests the purity of the isolated EVs and “cell-free” nature of the supernatant fractions **(Figure 1B and 1C)**. Consistent with the hypothesis that endolysosomal contents are released over the course of infection and process of CCV development, we observed an increase in the amount of the lysosome-associated membrane marker protein (LAMP1) and the pro-forms of the aspartyl protease cathepsin D, in the EV fraction of infected cells on 5 dpi, compared to that on 2 dpi **(Figure 1B and 1C)**. Cathepsin D is generated as pre-procathepsin D in the endoplasmic reticulum and transported through the trans-Golgi network to endocytic organelles. Procathepsin D is known to go through a two-step maturation process in the endolysosomal compartments: propeptide (∼52kDa) is partially cleaved in the first step to produce an active single-chain intermediate (∼48kDa), which is then further processed in the acidic lysosomal compartments to produce double chain, mature cathepsin D with the heavy chain ∼34kDa (31). Interestingly, we observed that infection was associated with a decrease in the levels of heavy chain form in the lysates, suggesting that *Coxiella* infection interferes with the maturation of cathepsin D, as has been demonstrated for cathepsin B in other cell types (23,32). The increase in levels of the pro-forms of cathepsin D in the EV fraction of infected cells on 5 dpi was correlated with a corresponding increase in the lysates, likely due to impaired processing to the mature forms. Cargo sorting and exosome biogenesis in MVBs occur via endosomal sorting complexes required for transport (ESCRT)-dependent and ESCRT-independent pathways, and consequently, exosomes may be identified by distinct molecular markers including the tetraspanin family proteins (CD63), ESCRT machinery proteins (Tsg101, Alix) (27–30,33). Notably, EV fraction was positive for Tsg101 and Alix (**Figure 1B and 1C**).

**Figure 1:**
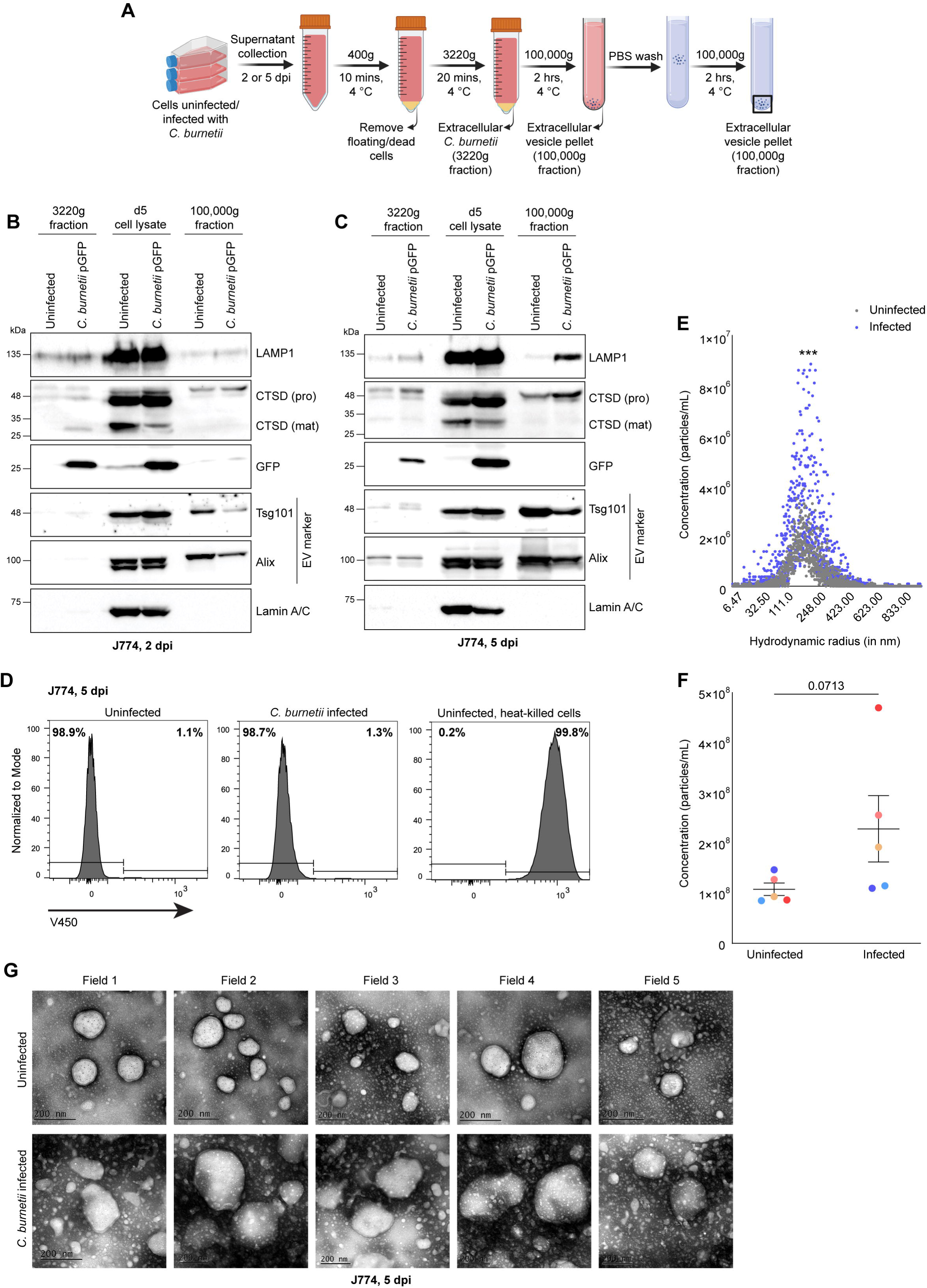
Infection with *C. burnetii* leads to increased detection of extracellular vesicles (EVs) and endolysosomal proteins in macrophages. J774 macrophages cells were either uninfected or infected with GFP-expressing *C. burnetii* (MOI 50) for 5 days. **(A)** Schematic of Extracellular vesicle (EV) isolation from conditioned media of J774 macrophages by differential centrifugation. **(B-C)** Cell-free supernatant was collected from uninfected or *C. burnetii*-infected cells on 2 dpi **(B)** and subsequently on 5 dpi, along with cell lysates **(C)**. Immunoblot of the 3220g fraction, lysates and 100,000g fraction derived from supernatants collected on 2 dpi **(B)** or 5 dpi **(C)** to detect various proteins, as indicated. **(D)** Cell viability was determined by flow cytometry by staining cells with LIVE/DEAD fixable violet dead cell stain on 5 dpi. J774 cells were heat-killed at 60°C for 20 minutes and were used as the positive control. Histogram shows the percentage of live cells (left) and dead cells (right) in the bulk population. One representative data set is shown. **(E)** Concentration of particles of different sizes in the EV fraction isolated from infected and uninfected macrophages by Nanoparticle Tracking Analysis. **(F)** Total concentration of particles in the EV fraction isolated from infected and uninfected macrophages with each dot representing a distinct experiment **(G)** Representative transmission electron microscopy images of the EV fraction isolated from uninfected or infected cells at 5 dpi (scale bars = 200 nm). Data presented is representative of at least 3 experiments **(B and C)**, 2 experiments **(D and G),** and assimilation of 5 experiments **(E, F)**.

An intense GFP band was observed in the lysates of cells infected with GFP-expressing *Coxiella*, indicating robust infection and intracellular bacterial replication **(Figure 1B and 1C)**. Concurrently, GFP was detected in the 3220g fraction on 2 as well as 5 dpi, indicating effective enrichment of extracellular bacteria in the 3220g fraction **(Figure 1B and 1C).** In order to determine whether the increase in extracellular levels of LAMP1 and cathepsin D was due to cell death, we determined cell viability on 5 dpi. Notably, *C. burnetii*-infected cells showed no prominent difference from that of uninfected cells, suggesting that the detection of endolysosomal proteins in the cell-free supernatants of the infected cells is independent of cell death **(Figure 1D)**. In order to assess any quantitative differences between the EV fractions of uninfected and infected cells, the EV fraction was analysed by nanoparticle tracking analysis (NTA) **(Figure 1E and 1F)**. The number of particles corresponding to each size was higher in the EVs from infected cells **(Figure 1E),** and the total concentration of particles (inclusive of all sizes) derived from infected cells, from 5 independent experiments, also showed a higher trend compared to that of uninfected cells **(Figure 1F)**. Negative staining transmission electron microscopy (TEM) validated the presence of intact vesicular particles in the EVs derived from uninfected as well as infected cells **(Figure 1G)**. Overall, these data suggest that *C. burnetii* infection augments exocytosis and extracellular vesicles and endolysosomal biomolecules are detected over the course of infection.

### Infection with *C. burnetii* enhances surface localization of LAMP1 in macrophages

Lysosomal exocytosis involves the fusion of LE/MVB/lysosomes with the plasma membrane (PM) to secrete luminal contents to the extracellular space (9). The enhanced detection of extracellular vesicles and cargo during *Coxiella* infection (**Figure 1**) prompted us to examine whether LE/lysosomal exocytosis may be involved by employing conventional and quantitative readouts of this process (16,34). Fusion of LAMP1+ compartments with the plasma membrane leads to detection of the luminal domain of LAMP1 on the cell surface (34). Having detected increased levels of LAMP1 in the EV fraction by immunoblotting in **Figure 1**, we examined the presence of cell-surface-associated LAMP1 (referred to as ‘sLAMP1’) on the plasma membrane by microscopy and flow cytometry. J774 macrophages left uninfected or infected with GFP-expressing *C. burnetii* were immunostained for LAMP1 (without permeabilization) at 5 dpi **(Figure 2A)**. Since exocytosis is highly regulated by intracellular calcium levels, Ionomycin, a calcium ionophore that promotes the influx of Ca^2+^ from extracellular media into the cells, was used as a positive control (35). The use of ionomycin allowed us to classify the observed surface localization of LAMP1 into three categories: no localization, partial localization, and total localization to allow accounting for a wider range of sLAMP1 localization patterns, as shown in the representative images **(Figure 2B)**. Using this binning strategy, we found that infected cells with a detectable vacuole containing GFP+ bacteria exhibited no or partial localization of sLAMP1 **(Figure 2C)**. Notably, the percentage of cells with detectable sLAMP1 was significantly higher in infected than in uninfected cells **(Figure 2C)**.

**Figure 2:**
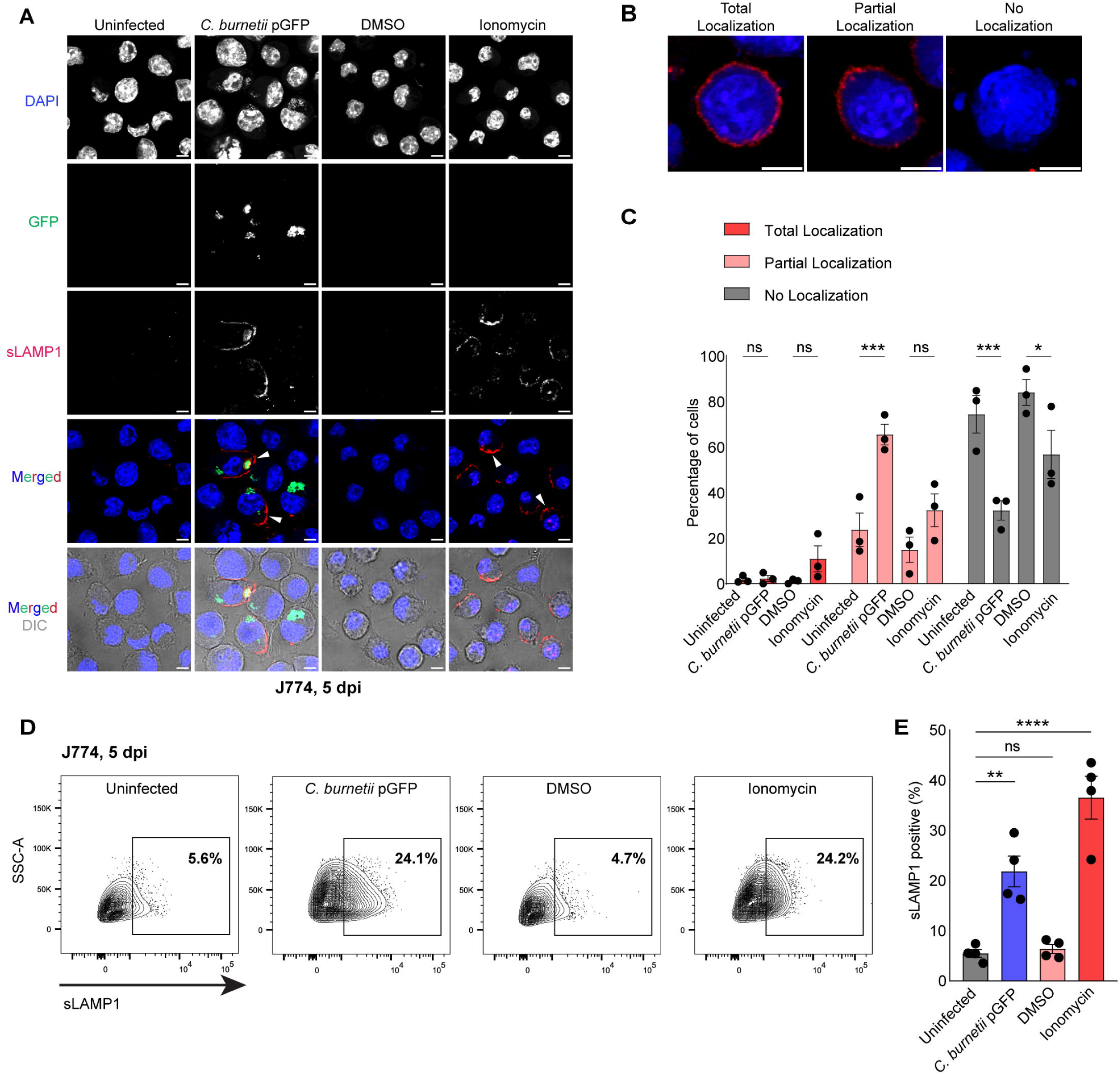
Infection with *C. burnetii* enhances surface localization of LAMP1 in macrophages. J774 macrophages were left uninfected or infected with GFP-expressing *C. burnetii* (MOI 50) for 5 dpi, or treated with ionomycin or vehicle control DMSO and stained to detect LAMP1, without permeabilization. Ionomycin was used as the positive control for exocytosis. **(A)** Immunofluorescent images of surface LAMP1 (sLAMP1) localization in J774 macrophages (scale bar indicates 10 µm). **(B)** Binning strategy used to categorize surface localization phenotypes as total, partial, and no localization, and is represented using ionomycin treatment (scale bars = 5 µm). **(C)** Quantification of sLAMP1 phenotypes under infection and ionomycin treatment conditions. Data are shown as the mean ± SEM of three independent experiments, with at least 90 cells quantified for each treatment, per experiment. One representative data set **(D)** and averages of four experiments **(E)** showing percentage of cells with sLAMP1 localization as determined by flow cytometry. For the *C. burnetii* pGFP-infected samples, sLAMP1+ cells were gated on the GFP-positive population.

These observations were further validated by measuring the percentage of sLAMP1+ cells by flow cytometry (**Figure 2D**). A significant increase in the percentage of sLAMP1+, *C. burnetii-*infected cells (gated for the cells infected with GFP-expressing *C. burnetii)* was noticed compared to that of uninfected cells, at 5 dpi, as shown in representative data (**Figure 2D**) or average across four independent experiments (**Figure 2E**). Taken together, these results indicate that infection with *C. burnetii* leads to increased fusion of LAMP1+ compartments with the PM, implicating LE/lysosomal exocytosis.

### Infection with *C. burnetii* increases the detection of extracellular endolysosomal proteins and sLAMP1 in non-phagocytic cells

To evaluate whether the infection-induced exocytosis and LAMP1 localization on the cell surface (sLAMP1) was common across different cell types permissive to *Coxiella,* we infected HeLa with GFP-expressing *C. burnetii* at an MOI of 100 for 6 days. The 3220g and 100,000g fractions were collected from the conditioned media using differential centrifugation as mentioned in **Figure 1A**. We observed an increase in the levels of LAMP1 and CD63 in the EV fraction of the infected cells at 6 dpi (**Figure 3A and B**). Consistent with our previous results, we detected Lamin A/C only in the lysates while the EV fraction of infected cells was relatively more abundant in exosome markers (Tsg101, Alix, CD63) and also microvesicle-associated markers (Annexin A1) (**Figure 3A and B**). Interestingly, we also detected some levels of GFP in the EV fraction. *Coxiella* is expected to be sedimented at 3220g, and hence it is less likely that these are intact bacteria. However, the possibility of residual bacteria, or smaller spore-like forms or GFP-protein associated with intracellular organelles co-sedimenting at 100,000g may be possible, particularly since there are no intermediate sedimentation steps between 3220g and 100,000g.

**Figure 3:**
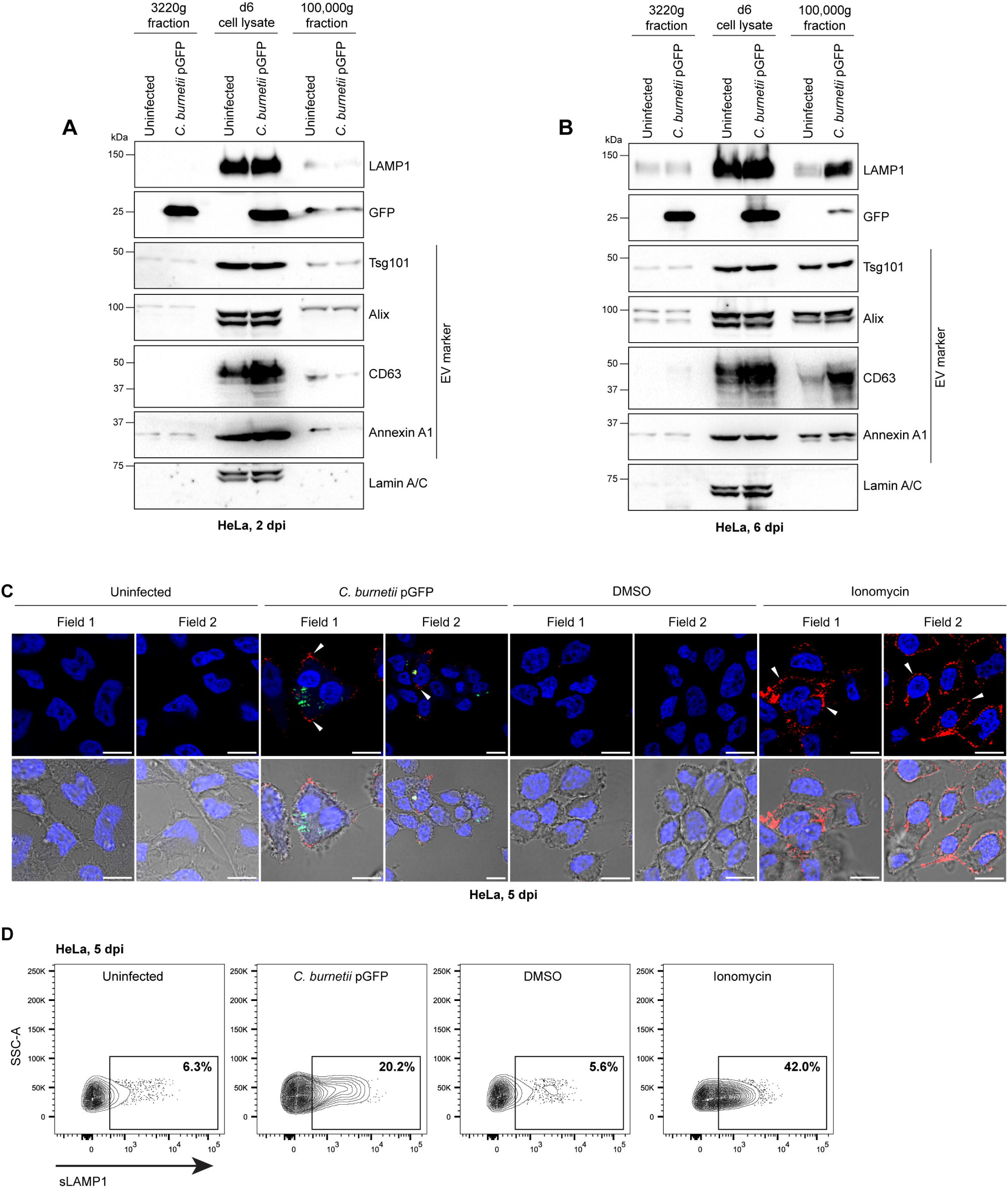
Infection with *C. burnetii* increases the detection of extracellular endolysosomal proteins and sLAMP1 in non-phagocytic cells (A-B) Immunoblot of HeLa uninfected or infected with GFP-expressing *C. burnetii* (MOI 100). Immunoblot of the 3220g fraction, and 100,000g fraction derived from cell-free supernatant collected from uninfected or *C. burnetii-*infected cells (MOI 100) on 2 dpi **(A)** and subsequently on 6 dpi, along with cell lysates **(B)**. **(C)** Immunofluorescent images of surface LAMP1 (sLAMP1) in HeLa left uninfected, infected with *C. burnetii* pGFP MOI 100, or treated with ionomycin or vehicle control DMSO and stained to detect LAMP1, without permeabilization. Scale bar indicates 10 µm. **(D)** One representative data set showing percentage of cells with sLAMP1 localization as determined by flow cytometry. For the *C. burnetii* pGFP-infected samples, sLAMP1+ cells were gated on the GFP-positive population. Data shown is representative of at least two experiments.

The levels of sLAMP1 were examined by confocal microscopy and flow cytometry as described for **Figure 2**. Consistent with J774, we detected LAMP1 on the surface of infected cells, and ionomycin treatment served as a positive control (**Figure 3C**). A similar increase in the percentage of sLAMP1+, *C. burnetii*-infected cells (gated for the cells with GFP fluorescence) compared to that of the uninfected was observed at 5 dpi, as examined by flow cytometry **(Figure 3D)**. Thus, an increase in exocytosis was observed in both *C. burnetii-*infected phagocytic as well as non-phagocytic cells.

### T4BSS activity of *C. burnetii* is essential for infection-induced exocytosis

Since infection-associated exocytosis was not likely due to cell death in our experimental settings **(Figure 1D)**, we examined the factors that regulate EV release during infection. To this end, *icmL*::Tn *C. burnetii* was used to infect J774 to determine if the activity of the *Coxiella* type IVB Dot/Icm secretion system (T4SS) and bacterial effectors was required. IcmL-deficient *C. burnetii* is impaired in the translocation of effectors and hence, replication inside host cells, as validated by the intracellular levels of the chaperone GroEL **(Figure 4A).** At 5 dpi, immunoblot analysis showed an increase in the amount of LAMP1 and pro-forms of cathepsin D in the EV fraction collected from wt-infected, but not the *icmL*::Tn-infected cells **(Figure 4A)**, indicating that T4BSS activity is required for exocytosis. In addition, the pro-forms of cathepsin D were higher in the lysates of wt *C. burnetii*-infected cells, but not those of untreated or *icmL*::Tn, with a corresponding reduction in mature cathepsin D levels in wt *C. burnetii*-infected cells, suggesting that the processing of procathepsin D is inhibited by T4SS activity. Thus, T4SS activity is required for interfering with the intracellular processing of procathepsin D and promoting the extracellular levels of LAMP1 and procathepsin D, in association with EV markers. No significant difference in the cell viability in wt or *icmL C. burnetii*::Tn-infected cells compared to the uninfected at 5 dpi was observed **(Figure 4B)**.

**Figure 4:**
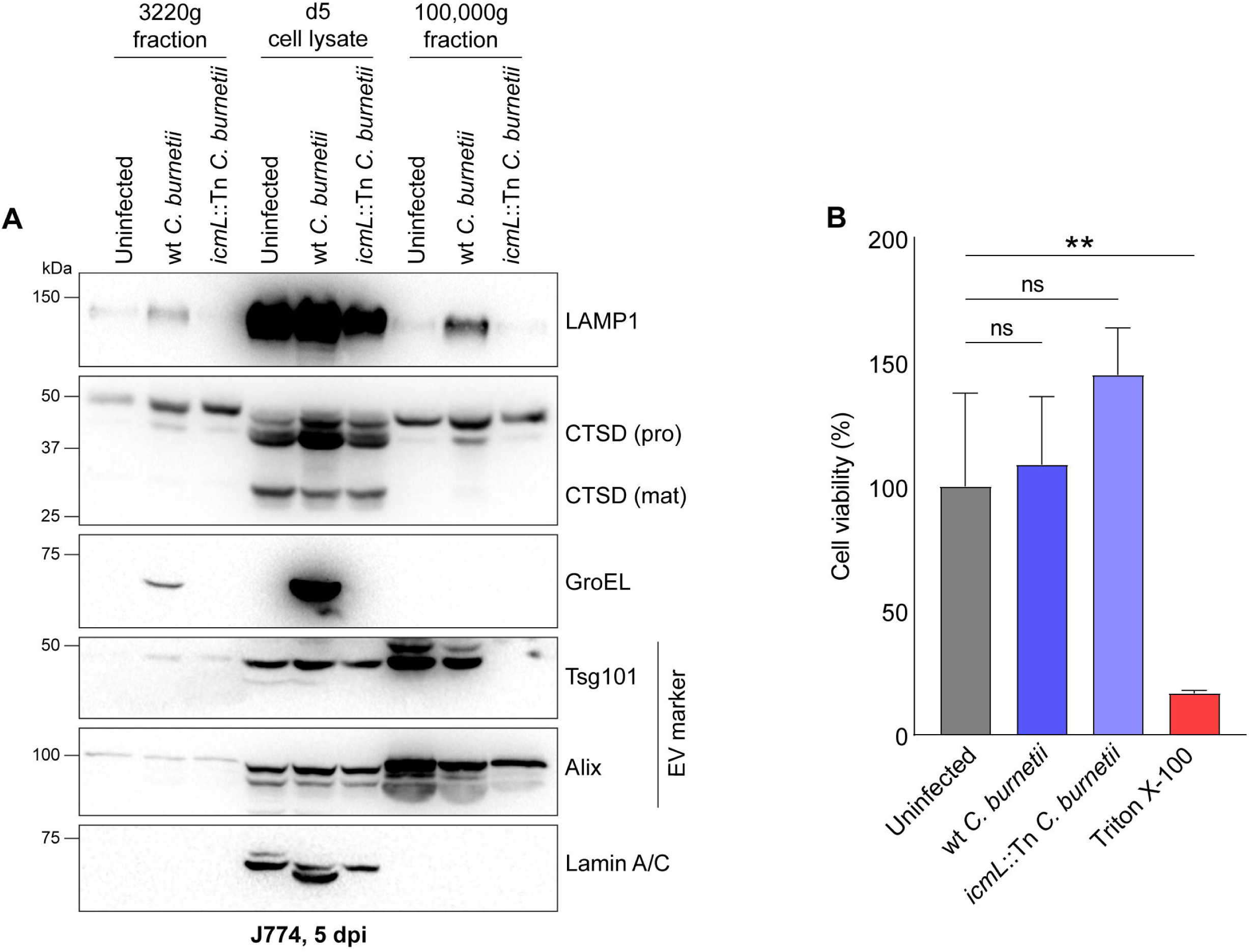
T4BSS activity of *C. burnetii* is essential for infection-induced exocytosis. **(A)** Immunoblot of J774 cells uninfected or infected with wt or *icmL*::Tn *C. burnetii* (MOI 50) at 5 dpi. Cell-free supernatant was collected from uninfected or *C. burnetii-*infected cells, along with cell lysates **(A)**. **(B)** Percentage of viable J774 cells by MTT assay at 5 dpi. Treatment with Triton X-100 is used as the positive control. Data presented is representative of at least 3 **(A)** or 2 independent experiments **(B)**.

### Modulation of the activity of the lysosomal cation channel mucolipin-1 (TRPML1) negatively impacts intracellular *C. burnetii* replication

To understand the host factors involved in regulating infection-associated exocytosis, we evaluated the role of the lysosomal divalent cation channel mucolipin-1 (ML-1/TRPML1), which controls TFEB activation in a Ca^2+^-dependent manner to trigger exocytosis (9,16,17). PI(3,5)P_2_ and PI(4,5)P_2_ have been reported to be the endogenous agonist and antagonist that regulate the activity of TRPML1 (36,37). Here, we used the commercially available ML-SA1 (mucolipin synthetic agonist 1), and ML-SI3 (mucolipin synthetic inhibitor 3) to modulate TRPML1 activity post-infection and evaluate the effect on bacterial replication (38–42). ML-SI3 has also been reported to inhibit the activity of ML-SA1-induced TRPML-1 activation (42,43). Treatment of infected cells with ML-SA1 (when added on 2 dpi) restricts intracellular bacterial replication in a dose-dependent manner, as observed with fluorescence expressed by *C. burnetii* pGFP **(Figure 5A),** without significant loss of cell viability **(Figure 5B)**. Interestingly, treatment with ML-SI3 also led to a decrease in bacterial replication **(Figure 5C)**, without ML-SI3 affecting cell viability **(Figure 5D).** Thus, treatment with either TRPML1 agonist or inhibitor decreases bacterial replication. These data suggest that either tight and/or temporal regulation of TRPML1 activity is required for optimal bacterial replication or that these synthetic agonist/antagonist skew TRPML1 activity to extremes, adversely affecting intracellular *Coxiella* replication. As the agonists or inhibitors used modulate not only TRPML1 activity, but also cellular levels of calcium as released by lysosomal organelles, the data are indicative that cellular calcium levels could play a nuanced role in *Coxiella* infection.

**Figure 5:**
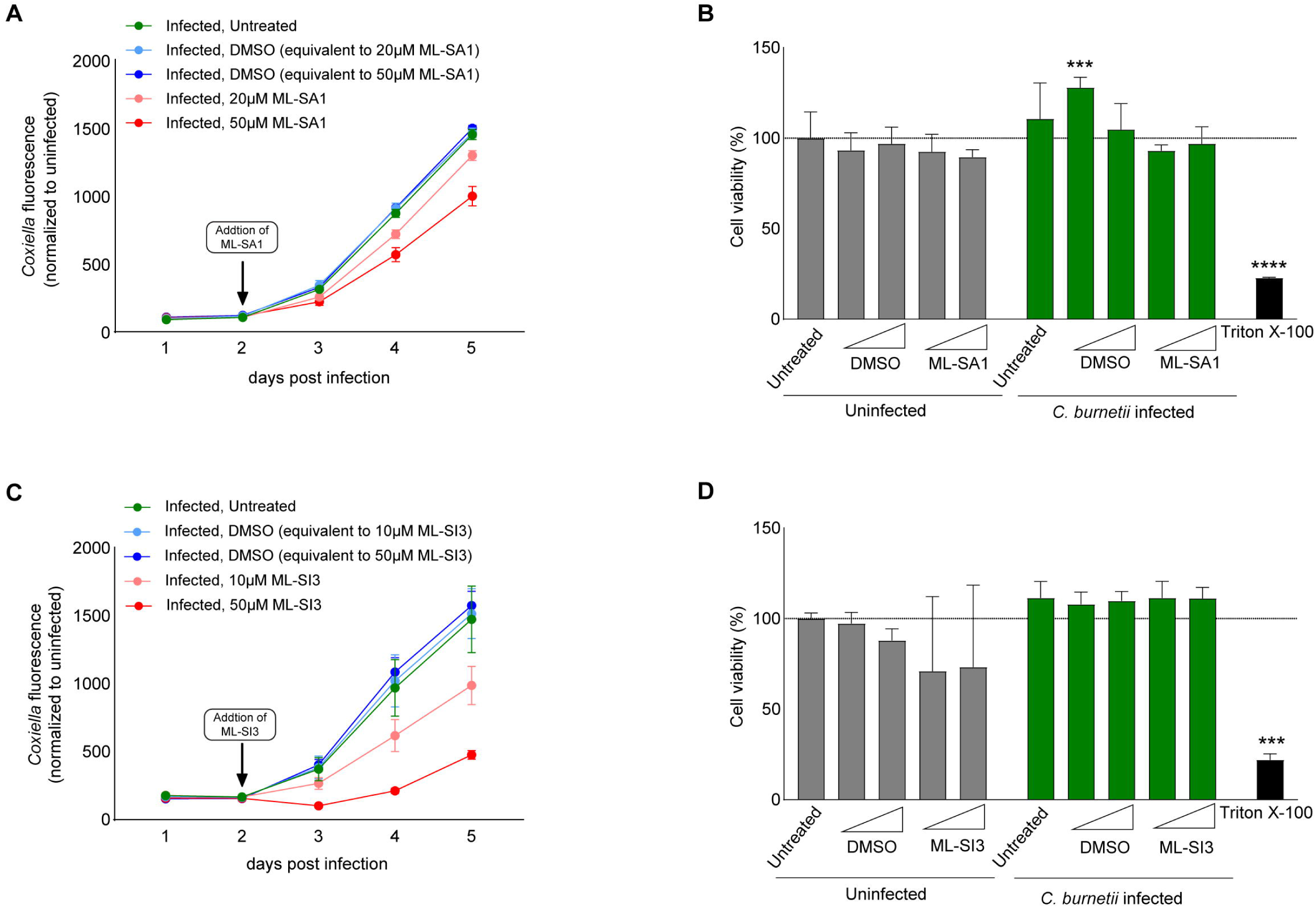
Modulation of the activity of the lysosomal cation channel mucolipin-1 (TRPML1) negatively impacts intracellular *C. burnetii* replication. HeLa cells were treated with different concentrations of the synthetic agonist of TRPML1, ML-SA1 **(A-B)** or inhibitor ML-SI3 **(C-D)** and intracellular replication and host cell viability was measured. Intracellular replication of *C. burnetii* pGFP was assessed by measuring GFP fluorescence **(A, C)**, and cell viability by MTT assay **(B, D)**. Data shown is representative of 3 independent experiments **(A, C)** and 2 experiments **(B, D)**.

## Discussion

This study characterizes *Coxiella burnetii* infection-induced exocytosis, its contribution to the intracellular replication of *C. burnetii,* and the host and bacterial factors that regulate this process. Data presented here demonstrates that infection enhances the detection of extracellular vesicles (EVs) in both phagocytic and non-phagocytic cells, as observed through (i) detection of endolysosomal proteins (LAMP1, cathepsin D), exosome associated markers including Tetraspanins (CD63), ESCRT proteins (Tsg101, Alix), and microvesicle-associated markers (Annexin A1) in the cell-free supernatants, and (ii) detection of LAMP1 on cell surface by indirect immunofluorescence as well as flow cytometry as shown in **Figures 1, 2 and 3**. TEM studies provided a microscopic validation of the presence of spheroid, extracellular vesicles in the EV fraction. While it is also possible that other vesicular extracellular carriers described as microvesicles, apoptotic bodies, blebbisomes, etc, are also released during infection, the average size observed by NTA **(Figure 1E)** is primarily indicative of exosomes. Notably, no significant cell death was observed at the time points where EVs were harvested, indicating that their release is not a consequence of compromised cell viability **(Figures 1 and 4)**.

Nanoparticle tracking analysis indicated that the total number of EVs and EVs of specific size ranges tend to be higher in those derived from infected cells **(Figure 1F)**. EVs collected from the infected cells also exhibited qualitative differences from those of uninfected cells, as determined by relative enrichment of proteins including LAMP1, procathepsin D and CD63 **(Figures 1, 3 and 4)**. Further, an enhanced secretion of the lysosomal protease procathepsin D in the EV fraction was observed, in correlation with decreased levels of processed, mature form in infected cell lysates and this effect was T4SS activity-dependent **(Figures 1C and 4A)**. These results are reminiscent of reduced processing and high secretion of procathepsin D and impaired lysosomal functions in HeLa cells infected with the T3SS SPI-2-sufficient *Salmonella* Typhimurium (44,45). At the time of preparation of this manuscript, Bird, L.E. et al. also reported that *C. burnetii* infection leads to secretion of lysosomal contents, including hydrolases, which aligns with our observations (32).

Infection-induced exocytosis was observed to require concerted functions of bacterial and host factors. Extracellular LAMP1 levels were absent when cells were infected with *icmL*::Tn *C. burnetii,* suggesting that bacterial T4SS activity is required **(Figure 4A)**. Since the T4SS-deficiency impairs not only the translocation of bacterial effectors into host cells but also the generation of a replication permissive, fusogenic vacuole, decreased exocytosis may be attributed to lack of either specific effector functions or impairment of maturation/progression of the CCV to a stage that is positioned for interception and engagement with the host exocytic pathways.

Inhibiting TRPML1 function by ML-SI3 or activating the same using the synthetic agonist ML-SA1 has an inhibitory effect on *Coxiella* replication **(Figures 5A and C)**. Interestingly, this observation is corroborated by a recent study demonstrating that MLSA-5 (another TRPML1 agonist) decreases *Coxiella* replication and CCV biogenesis (25). It is possible that agonist-induced activation of TRPML1 on 2 dpi (as in **Figure 5A**) causes excessive, premature, or untimely exocytosis and interferes with the early development of the CCV. Indeed, *Coxiella* employs effectors such as Vice (CBU_2007) and CvpE to facilitate early CCV expansion by the induction of macropinocytosis, inhibiting ESCRT pathways, and interfering with tubulation of lysosome-like vacuoles (25,46). Further, CvpE was demonstrated to disrupt PIKfyve activity, interfering with the synthesis of endogenous TRPML1 agonist PI(3,5)P2 and TRPML1 activity (25). Together, these results show that TRPML1 activity is tightly regulated for optimal bacterial replication during infection.

Late-stage induction of apoptosis (∼7 dpi) was recently reported to facilitate bacterial egress in infected human endothelial cells (47). The experimental approach presented here also allowed us to assess extracellular bacteria, as part of the ‘cell-free, nucleus-free’ 3220g fraction, and factors affecting potential exit of bacteria from infected cells over the course of infection. Bacterial-expressed GFP or GroEL was observed in the 3220g fraction derived from wt *C. burnetii-infected* cells, but not from *icmL*::Tn *C. burnetii*-infected cells, indicating that these are newly formed bacteria over the course of the infection cycle **(Figures 1, 3 and 4)**. Concurrently, LAMP1 is also detected in the 3220g fraction by 5-6 dpi, in addition to *Coxiella*, raising the possibility that *Coxiella* is released from infected cells, at least in part, in membrane-enclosed compartments. Indeed, Schulze-Luehrmann, J. et al. also observed events indicating fission of CCV, bacterial exit without death of infected host cells, and bacterial egress to be non-synchronous (47), which adds support to this hypothesis. We speculate that, in addition to apoptotic death-associated release of free *Coxiella*, multiple mechanisms may contribute to bacterial egress, including *Coxiella* release in membrane-bound vesicles.

Altogether, the data presented here, using *Coxiella burnetii* as a model system, demonstrate that infection with a lysosomal vacuolar pathogen promotes the release of heterogeneous extravesicular populations over the course of infection. Of note, the secretory pathway, cargo traffic between endosomes and ER-Golgi, all contribute to CCV biogenesis, implying the pathogen’s ability to subvert multiple host vesicle traffic pathways (32,48–50). There is growing evidence for how intracellular pathogens subvert host cell exocytosis for their entry, sustained replication, and dissemination from infected cells (51–53). *Listeria monocytogenes* subverts exocytosis, host exocyst complex to promote membrane protrusions and exit from infected cells (52,54). The obligate intracellular vacuolar bacterial pathogen, *Chlamydia trachomatis,* was demonstrated to exit from infected cells as free *Chlamydia* by lysis of the inclusion vacuole and the infected cell or extrusion of parts of the bacteria-containing vacuole in membrane-bound protrusions without inducing host cell death (55). *Anaplasma phagocytophilum,* which replicates in MVB-like compartments, can exit host cells by co-opting the MVB exocytosis (56). Of note, CCV also displays many MVB-like characteristics, including LBPA-rich membrane and fusogenicity (46). Exocytosis also facilitates non-lytic release of exosome-enclosed uropathogenic *E. coli* (UPEC) from bladder epithelial cells, where infection-mediated neutralization of pH of UPEC-containing lysosomal vacuoles induces the activity of TRPML3 and thereby, exocytosis (57). Notably, *Coxiella* infection has also been demonstrated to reduce acidification of lysosomes-derived CCV and maintain it at pH ∼5.2 compared to that of conventional lysosomes (∼4.8) (23). In the case of *Coxiella* infections, the precise physiological relevance of exocytosis, extruded vesicles and cargo, cell-to-cell communication, release of PAMPs/DAMPs, their implications for bacterial dissemination, and the bacterial effectors that potentially regulate exocytosis require extensive investigation and are exciting research directions for the future.

## Acknowledgements

The support for this work and associated project personnel (KB) was provided by the start-up research grant (SRG) from the Department of Science and Technology-Science and Engineering Research Board, DST-SERB (now Anusandhan National Research Foundation, ANRF), awarded to SG (SRG/2022/002157). Our work was additionally supported by Indian Institute of Science Education and Research Thiruvananthapuram (IISER TVM) and DBT/Wellcome Trust India Alliance Intermediate Fellowship awarded to SG (IA/I/23/2/507001). AB is supported by the Department of Biotechnology-Junior Research Fellowship (DBT-JRF, DBT/2022-23/IISER-TVM/2125) and HK and NS by IISER TVM. We extend our gratitude to Prof. Craig Roy (Yale University School of Medicine) for generously providing the *Coxiella burnetii* strains used in this study. We thank Dr. Karthik Chandiran (IISER TVM) for guidance with flow cytometry and data analysis and Dr. Karthik Subramanian (RGCB Thiruvananthapuram) for guidance with NTA, and optimization of EV staining procedures for TEM. We are thankful to Prof. Subba Rao Gangi Setty (IISc Bangalore) and Dr. Anshul Bhatt (Prof. Subba Rao Gangi Setty research group), for their insightful suggestions during the optimization of calcium-based exocytosis assays and Prof. Dipshikha Chakravortty (IISc Bangalore) for valuable feedback. We are grateful for the infrastructural support provided by the Biophysical Instrumentation Facility, Microscopy Core, Flow Cytometry Core and various labs within the School of Biology, and the Central Instrumentation Facility, IISER TVM. We thank the Centre for Biomaterials Cellular and Molecular Theranostics Testing Laboratories (CBCMTL), Vellore Institute of Technology, for NTA. We acknowledge Biorender for providing a platform for creating the figures.

## Materials and Methods

### Antibodies and Reagents

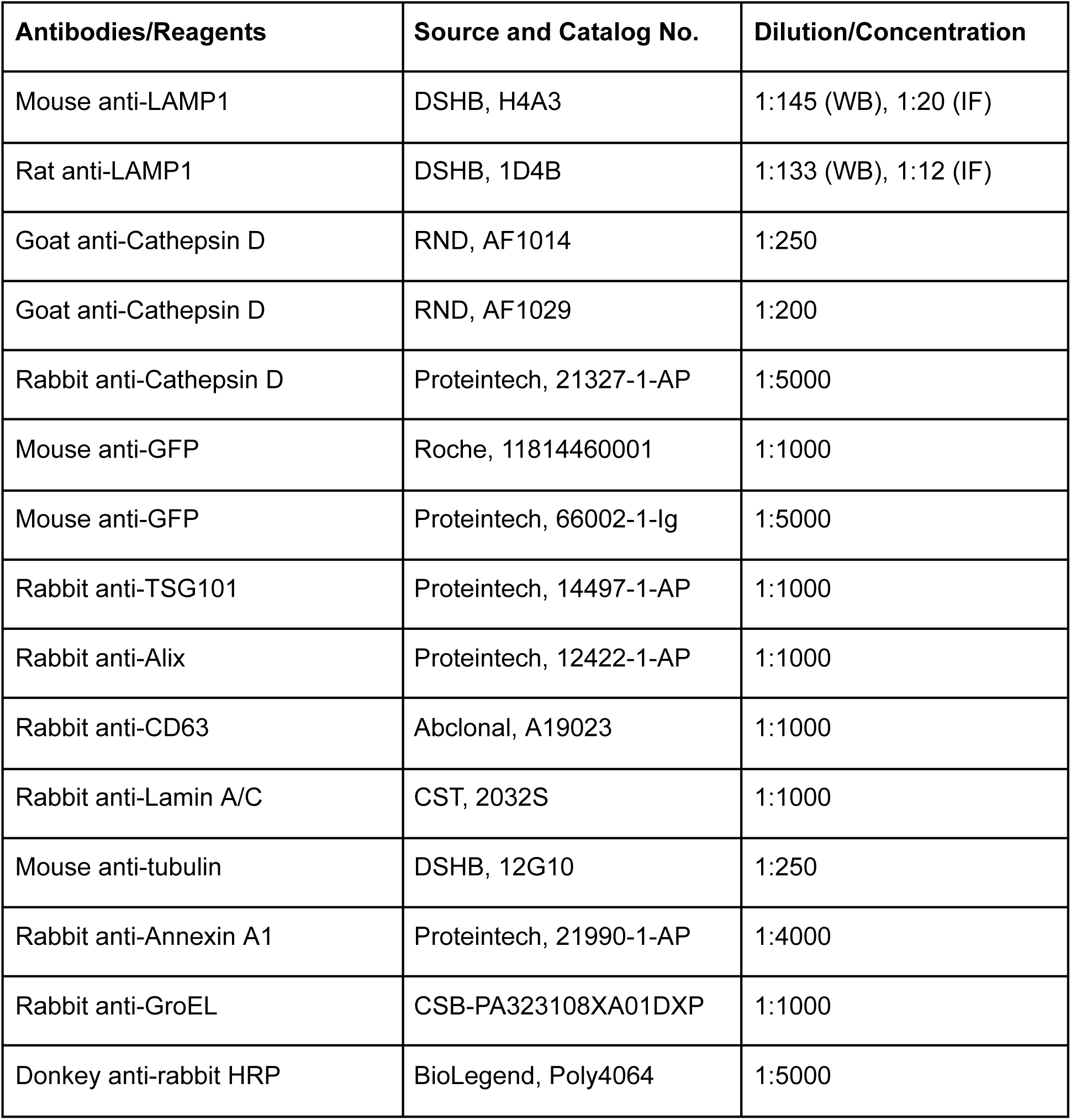
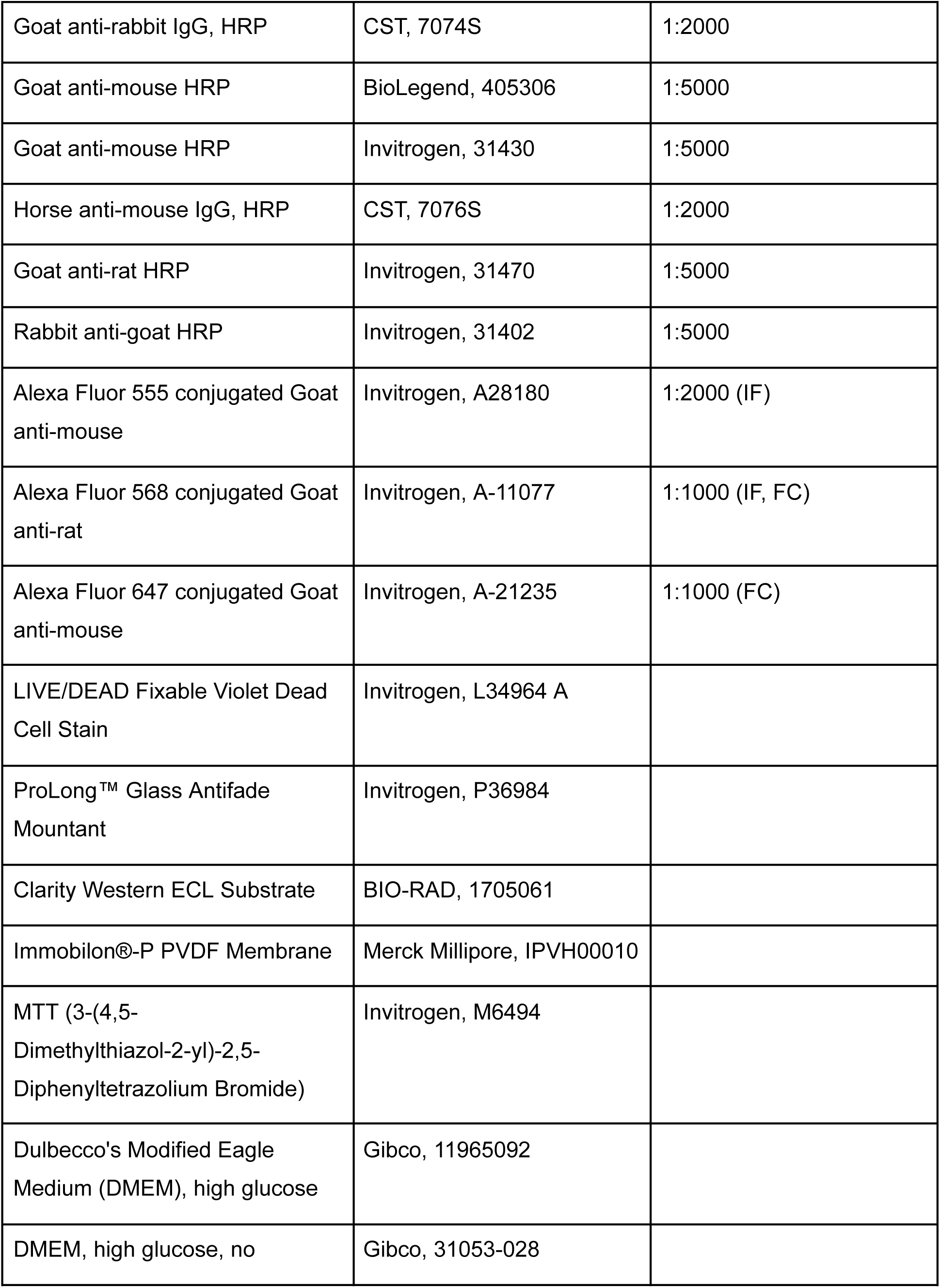
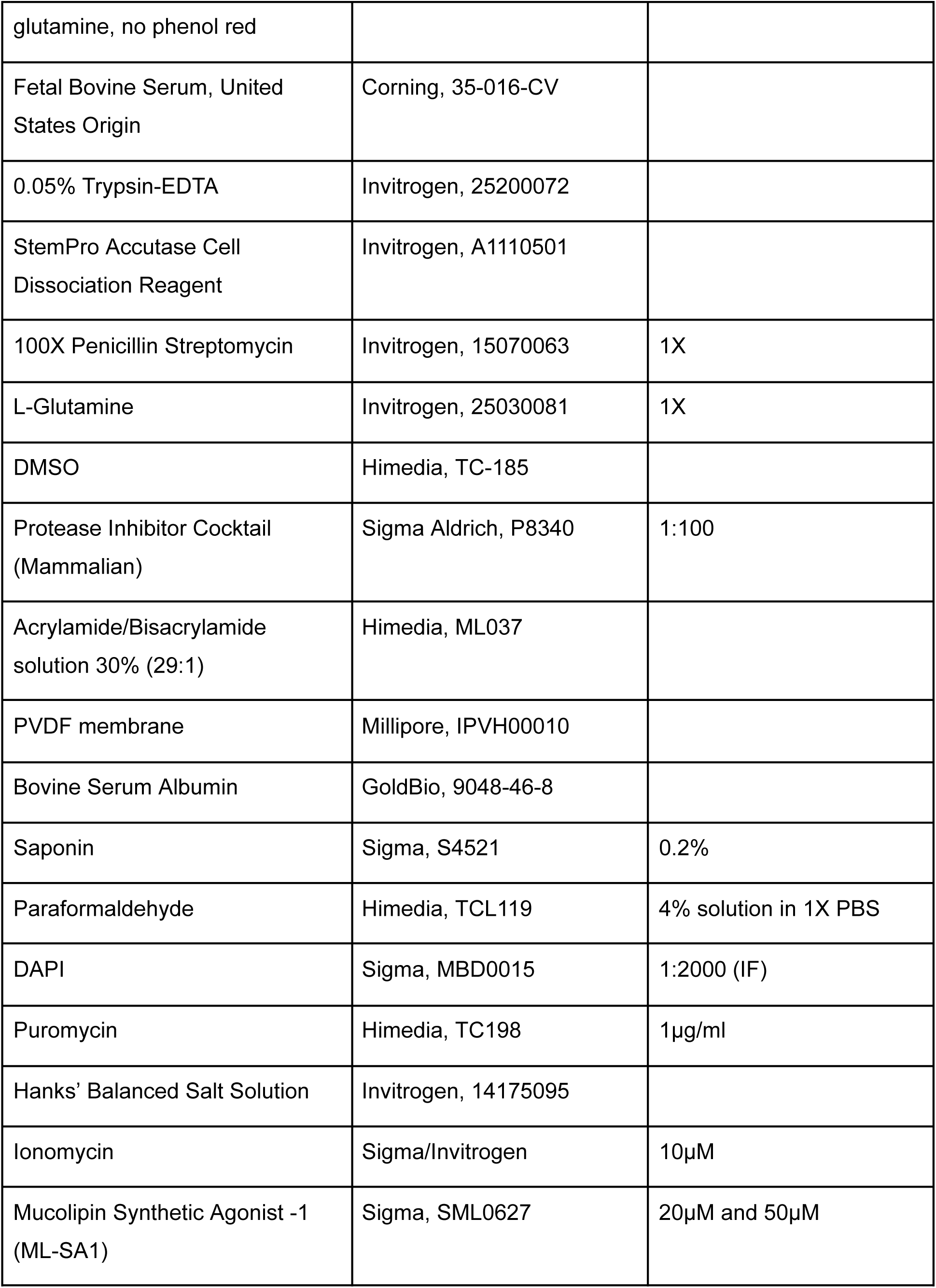
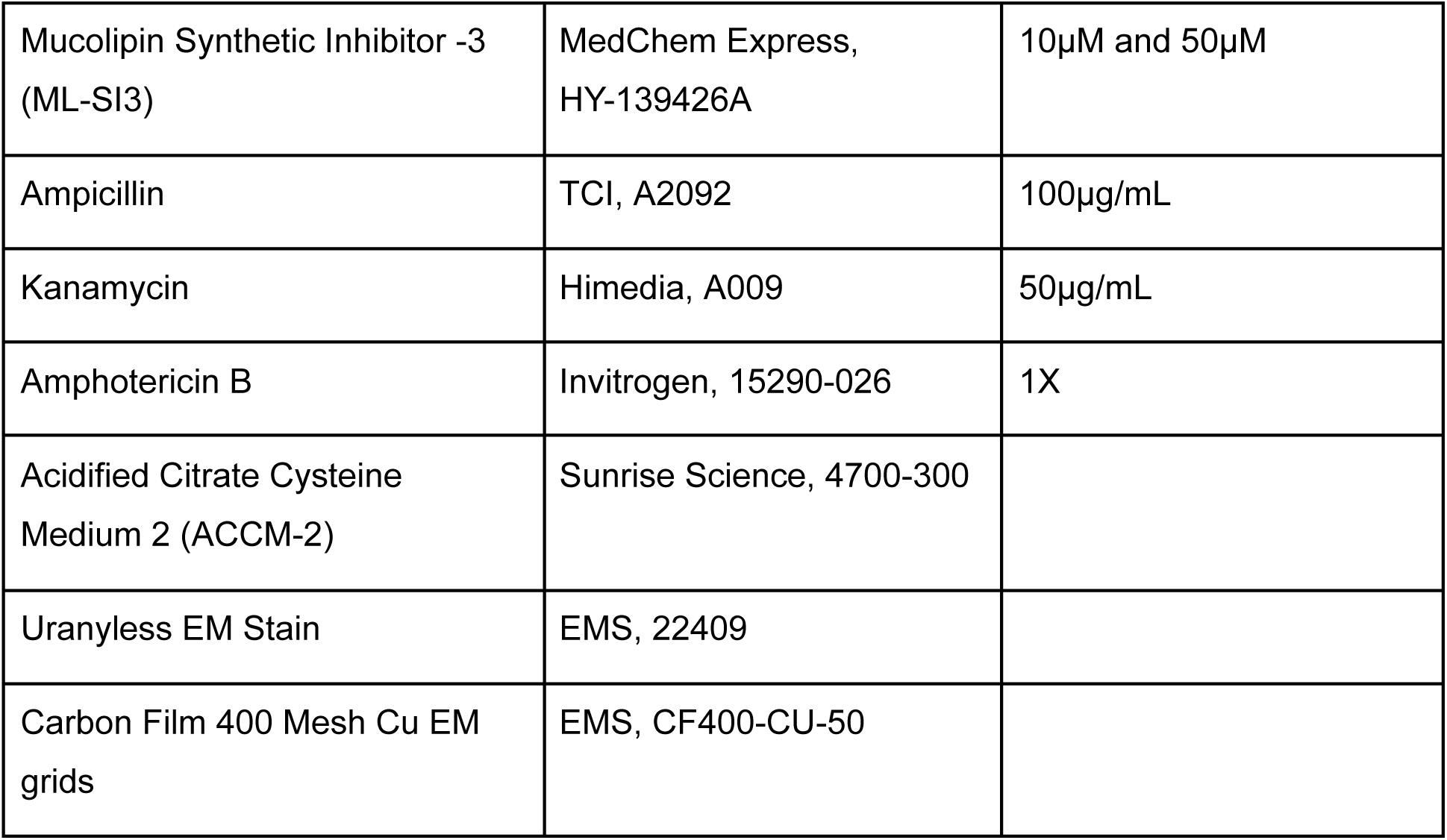

### Cell lines and cell culture

HeLa CCL2 and J774 cells were cultured in DMEM media supplemented with 10% fetal bovine serum (FBS), subsequently referred to as complete medium, in a humidified cell culture incubator with 5% CO_2_ at 37°C. For HeLa-WT, 1X penicillin/streptomycin and 1X amphotericin B were additionally added to the media. For the EV isolation, imaging, and flow cytometry experiments, cells were cultured in DMEM supplemented with 5% FBS. Cells were cultured in phenol red-free media supplemented with 10% FBS and 1% L-Glutamine for the plate reader-based growth curve experiments.

### C. burnetii strains

*C. burnetii* strains were grown for 6 days in ACCM-2 at 37°C, 2.5% O_2,_ and 5% CO_2_. Bacterial cultures were then centrifuged at 3220 g for 15 minutes at 4°C. The bacterial pellet obtained was resuspended in DMEM supplemented with 5% FBS. Genomic equivalents quantification was done by qPCR using *C. burnetii-*specific dotA primers (dotA F: GCGCAATACGCTCAATCACA, dotA R: CCATGGCCCCAATTCTCTT).

**Table 1:** *C. burnetii* strains used in this study.

| Sl. No | Name of the strain | Property | Source |
| --- | --- | --- | --- |
| 1. | wt *C. burnetii* | Wild type *C. burnetii* Nine Mile phase II (NMII) (RSA439) | Prof. Craig Roy, Yale University School of Medicine (7,58) |
| 2. | *C. burnetii* pGFP | *C. burnetii* that constitutively expresses GFP |  |
| 3. | *icmL::Tn C. burnetii* | *C. burnetii* with a transposon insertion in *icmL.1/dotI* (cbu1629) |  |

### Isolation of EVs

For EV isolation, J774 and HeLa cells were plated on 10 cm dishes (2 million/dish and 1.5 million/dish, respectively) and infected with *C. burnetii* (wt, or GFP-expressing, as indicated). For each experimental condition, 2-4 dishes were maintained. Cell supernatants were collected on indicated days, and cell debris was removed by centrifugation at 400 × g, 10 minutes, 4°C. Pellets were discarded, and supernatants were then spun at 3220 × g, 20 minutes, 4°C to obtain extracellular *Coxiella*. Pellets were stored at −20°C, and supernatants were ultracentrifuged (Type 50.2 Ti or SW40 Ti Rotor, Beckman Coulter) at 100,000 × g, 2 hours, 4°C to obtain small EVs. Pellets were washed in phosphate-buffered saline (PBS) 1X PBS and re-pelleted at 100,000 × g, 2 hours, 4°C. The pellets were resuspended in 20μl 1X filter-sterile PBS and stored for further processing.

### Western blotting

For the preparation of lysates, cells were washed with 10 mL 1X PBS after removal of supernatants on 5 dpi (J774) or 6 dpi (HeLa), and centrifuged at 400 × g, 10 minutes, 4°C. Cells were lysed in RIPA buffer (1% NP-40, 1 mM EGTA, 2 mM MgCl_2_, 0.5% Na^+^ Deoxycholate, 0.1% SDS, 150 mM NaCl, 50 mM Tris pH 7.5) supplemented with Protease Inhibitor Cocktail at 1:100 dilution on ice for 20 minutes with intermittent vortexing. The samples were centrifuged at 14,000 × g for 10 minutes at 4°C. Resultant lysates, 3220g fraction, and EV fraction were mixed with 4X Laemmli buffer supplemented with β-mercaptoethanol and boiled at 99°C for 5 minutes. Cell lysates were loaded in the ratio of 1:30 (J774 macrophages) and 1:60 (HeLa CCL2). Proteins were separated by 9% acrylamide/bisacrylamide gel electrophoresis, transferred to PVDF membranes, and probed with the indicated primary antibody, followed by horseradish peroxidase (HRP)-conjugated secondary antibody. Clarity Western ECL Substrate was used for chemiluminescence, and membranes were imaged using Bio-Rad ChemiDoc XRS+ system.

### Nanoparticle Tracking Analysis

1 μL of EV fraction isolated from uninfected and infected J774 macrophages was diluted to 1000μL using 1X filter-sterilized PBS and analyzed in a Particle Metrix Zetaview-PMX 130-Mono laser for measuring the concentration and size of particles at the Centre for Biomaterials Cellular and Molecular Theranostics Testing Laboratories (CBCMTL), Vellore Institute of Technology.

### Transmission electron microscopy

Negative staining was done for TEM sample preparation. Carbon film-coated 400 mesh copper EM grids were glow-discharged for 5 minutes using a GloQube® Plus glow discharge unit (Quorum Technologies). 10μL of the diluted EV suspension (1:10) was placed on a parafilm, above which the copper grid was placed for adsorption overnight at 4°C. After incubation, the samples were washed twice with 1X PBS and blotted with Whatman no. 1 filter paper. The EV sample was then fixed with 5μL of 2% glutaraldehyde for 30 seconds. This was followed by adding 10μL Uranyless solution on the surface of the EM grid for 1 minute, followed by blotting using filter paper. Next, the grid was allowed to air dry for 10 minutes and stored in an EM grid box. This grid was then visualized using a 120kV FEI Tecnai Spirit Bio-Twin Transmission Electron Microscope.

### Ca^2+^-induced lysosomal exocytosis assay

Ca^2+^-induced lysosomal exocytosis assays were performed as described previously by Escrevente et al. with slight modifications (35). Lysosomal exocytosis was induced using ionomycin. HeLa and J774 cells were incubated in Ca^2+^- and Mg^2+^-free ice-cold Hanks Balanced Salt Solution (HBSS) with DMSO or 10µM ionomycin in the presence of 4 mM CaCl_2_, for 10 minutes at 37°C. After completion of incubation, cells were placed on ice and processed for microscopy or flow cytometry analysis.

### Immunofluorescence

For immunofluorescence microscopy, J774 and HeLa cells were seeded on poly-L-Lysine-coated coverslips in 24-well plates, and left uninfected or infected with GFP-expressing *C. burnetii* at MOI 50 and 100, respectively. Post-infection, the plates were centrifuged at 500 x g for 30 minutes. On day 5 post-infection, ionomycin was used as a positive control, and cells were immunostained. For sLAMP1 localization, cells were washed three times with 1X PBS at desired time points, followed by blocking in a solution containing 0.5% BSA and 1% FBS in 1X PBS. Coverslips were then incubated with anti-LAMP1 primary antibody in blocking solution for 30 minutes at 4°C, washed thrice with 1X PBS, and fixed with 4% PFA in 1X PBS for 10 minutes. Post-incubation coverslips were washed with 1X PBS. Subsequently, coverslips were incubated with Alexa Fluor-conjugated secondary antibody and DAPI in a blocking solution for 1 hour at room temperature. Finally, the coverslips were washed thrice with 1X PBS and mounted with ProLong™ Glass Antifade Mountant.

Images were acquired on an Olympus confocal microscope (FluoView3000) using a 100×/1.35 NA oil immersion objective. Images were processed with Fiji software.

### Flow cytometry

For sLAMP1 analysis, J774 macrophages and HeLa CCL2 cells were seeded in 12-well (150,000 cells/well) and 24-well plates (25,000 cells/well) and left uninfected or infected with GFP-expressing *C. burnetii* at MOI 50 and 100, respectively. On day 5 post-infection, Ionomycin was used as a positive control, and cells were harvested using Accutase and blocked in the flow cytometry buffer (1X PBS + 2% FBS) for 30 minutes at 4°C. Cells were stained with anti-LAMP1 primary antibody for 30 minutes at 4°C. Next, cells were washed with 1X PBS and fixed using 4% PFA for 10 minutes at 4°C. Cells were again washed with 1X PBS and stained for Alexa Fluor 568 or Alexa Fluor 647 conjugated secondary antibody for 1 hour at RT. Finally, cells were washed with 1X PBS and resuspended in the flow cytometry buffer. Acquisition was performed in a BD FACSLyric^TM^ or BD FACSAriaIII flow cytometer. For the infected samples, we gated additionally on GFP-positive cells and then evaluated the sLAMP1 intensity. The percentage of sLAMP1-positive cells was determined using FlowJo software (BD Biosciences).

For cell viability analysis, J774 macrophages were left uninfected or infected with GFP-expressing *C. burnetii* at an MOI of 50. Cells were harvested 5 days post-infection using Accutase. Next, cells were resuspended in LIVE/DEAD Fixable Violet Dead Cell Stain as per the manufacturer’s instructions. For the positive control sample, cells were heat-killed by placing them at 60°C for 20 minutes before staining. Next, cells were washed with 1X PBS and fixed using 4% PFA for 10 minutes at 4°C. Finally, cells were washed with 1X PBS and resuspended in the flow cytometry buffer. Acquisition was performed in a BD FACSLyric^TM^. The data was plotted using FlowJo software (BD Biosciences).

### Bacterial growth curve experiments and MTT assay

Cells were seeded in 96-well black tissue culture plates with transparent bottom and infected on the next day with *C. burnetii* pGFP typically at multiple MOIs, including MOIs of 50, 100 and 250. Following infection, the plates were centrifuged at 500 x g for 30 minutes. From day 1 post-infection till day 5 post-infection, the fluorescence was measured in the Infinite F200 Pro Tecan plate reader at Ex/Em 485/538 nm. For experiments with small molecules, at two days post-infection, cells were treated with either of the two different concentrations of ML-SA1 (20μM and 50μM) or ML-SI3 (10μM and 50μM), or a corresponding volume of DMSO, and the data represented was normalized to uninfected.

After 5 days post-infection, MTT (5mg/ml) dissolved in 1X PBS was added to the wells after removing the media. Treatment with Triton X-100 for 30 minutes at 37°C was used as a positive control. After 4 hours of incubation at 37°C, DMSO was added to the wells and mixed well. Absorbance was measured using Infinite F200 Pro Tecan plate reader at 540 nm.

### Statistical analysis

The data were analyzed using GraphPad Prism 10.4.1 software. The statistical significance was calculated using different analyses: Mann-Whitney U test, t-test, one-way or two-way analysis of variance (ANOVA) with SD or SEM and post-hoc tests, as applicable to the data. Statistical differences were highlighted as * p ≤ 0.05, ** p ≤ 0.01, *** p ≤ 0.001, **** p ≤ 0.0001.

## Notes

### Competing Interest Statement

The authors have declared no competing interest.

